# Systematic differences between visually-relevant global and local image statistics of brain MRI and natural scenes

**DOI:** 10.1101/271338

**Authors:** Yueyang Xu, Ashish Raj, Jonathan Victor, for the Alzheimer’s Disease Neuroimaging Initiative

**Affiliations:** Stanford University Stanford, CA 94305; Department of Radiology Weill Cornell Medical College New York, NY 10065; Brain and Mind Research Institute and Department of Neurology Weill Cornell Medical College New York, NY 10065

**Keywords:** magnetic resonance imaging, brain, image statistics, human vision

## Abstract

An important heuristic in developing image processing technologies is to mimic the computational strategies used by humans. Relevant to this, recent studies have shown that the human brain’s processing strategy is closely matched to the characteristics of natural scenes, both in terms of global and local image statistics. However, structural MRI images and natural scenes have fundamental differences: the former are two-dimensional sections through a volume, the latter are projections. MRI image formation is also radically different from natural image formation, involving acquisition in Fourier space, followed by several filtering and processing steps that all have the potential to alter image statistics. As a consequence, aspects of the human visual system that are finely-tuned to processing natural scenes may not be equally well-suited for MRI images, and identification of the differences between MRI images and natural scenes may lead to improved machine analysis of MRI.

With these considerations in mind, we analyzed spectra and local image statistics of MRI images in several databases including T1 and FLAIR sequence types and of simulated MRI images,[1]–[6] and compared this analysis to a parallel analysis of natural images[7] and visual sensitivity[7][8]. We found substantial differences between the statistical features of MRI images and natural images. Power spectra of MRI images had a steeper slope than that of natural images, indicating a lack of scale invariance. Independent of this, local image statistics of MRI and natural images differed: compared to natural images, MRI images had smaller variations in their local two-point statistics and larger variations in their local three-point statistics – to which the human visual system is relatively insensitive. Our findings were consistent across MRI databases and simulated MRI images, suggesting that they result from brain geometry at the scale of MRI resolution, rather than characteristics of specific imaging and reconstruction methods.

## 1 Introduction

Development of image processing systems is often guided by the strategies used by the human visual system. The basic reason for this approach is that drawing inferences from images is a complex and ill posed problem, but one that the human visual system, as a result of evolutionary and developmental forces, has become reasonably effective at solving.

What is unclear, however, is the level of detail at which human vision should be taken to provide useful guidance: not only does it operate under different constraints, but also, it is matched to the statistical properties of images that result from projections of the natural environment onto the retina. This matching is at a surprising degree of detail – including not only the well-recognized stage of redundancy reduction by removal of global correlations [9], [10], but also an analysis of local statistics in a way that is closely matched to the features that distinguish patches of natural images [7]. While the computational strategies in human vision are sufficiently robust and general to enable perceptual judgments about images that are highly non-natural – for example, modern art – there is ample evidence that these strategies reflect a specific allocation of computational resources: for example, some kinds of local correlations are readily perceptible, while others, which are of comparable mathematical complexity, escape our notice [11].

Therefore, understanding the extent to which a particular image class shares the statistical properties of natural images is likely to be helpful in translating the lessons learned by the human visual system into specific applications. Here, we focus on this comparison for brain MRI.

Because the human visual system is matched to natural images both in terms of their global and local statistical properties, we consider both of these aspects of MRI images independently. Specifically, we analyze global statistical properties via the spatial power spectrum, and we analyze local statistical properties in terms of multipoint correlations. The latter analysis is carried out on whitened images, so it is independent of any overall differences in the power spectrum.

We may anticipate differences both in terms of power spectra and multipoint correlations. In principle, differences in power spectra may arise because natural scenes are equally likely to be viewed from many distances and angles but MRI images have a stereotyped size and framing. Thus, the spectral scaling behavior associated with natural scenes may not be shared by MRI. Differences in multipoint correlations may arise because of the differences in the imaging process itself. For example, in natural images, local features such as “T-junctions” arise because of occlusion [12]. But occlusion is not relevant to tomographic reconstruction, so T-junctions, when present, are likely to indicate a meeting-point of three objects. In addition, MRI acquisition involves computational reconstruction, and the details of this multistage process will influence the statistics of the image that is presented to the observer.

As we describe, we find substantial differences in both global and local statistics between MRI images and natural scenes. For both kinds of statistics, many of these differences are consistent across T1 weighted and FLAIR datasets. These findings suggest several paths for development of novel machine vision tools, including extension of existing algorithms through recognition of the distinctive statistical features of MRI images and the development of new classes of algorithms to mitigate the mismatch between MRI images and the images our visual system has evolved to process.

## 2 Materials and Methods

### 2.1 Databases and Image Selection

Analyses were carried out on de-identified human brain MRI images obtained from three databases and on simulated MRI images computed by BrainWeb [1]–[4]. The human MRI databases were: the Open Access Series of Imaging Studies (OASIS) [5], the Alzheimer Disease Neuroimaging Initiative (ADNI) [6], and a dataset from healthy volunteers and individuals diagnosed with a variety of neuroinflammatory diseases (mostly multiple sclerosis) collected in the Translational Neuroradiology Section of NINDS (here designated the TNS dataset). The TNS dataset was provided by Daniel S. Reich at the NINDS. Characteristics of the databases are summarized in Table 1.

All images were analyzed as sagittal slices, with a voxel size of 1.0 × 1.0 × 1.0 mm for the ADNI and TNS databases, and 1.0 × 1.0 × 1.25 mm for the OASIS database. For the OASIS and ADNI databases, images were obtained using T1weighted sequences[5][6]. For the TNS database, images were obtained with a T2-weighted FLAIR sequence.

The OASIS database, 380 datasets, consists of healthy subjects and patients with a variety of diagnoses, age range 18 to 96 years. The ADNI database, 119 datasets, consists of patients diagnosed with Alzheimer’s disease, age range 55 to 90 years. The TNS database had two subsets: one with 39 healthy subjects and one with 255 patients suspected of having neuroinflammatory diseases, such as multiple sclerosis; combined age range was 18 to 83 years. All databases included male and female subjects, and all datasets were analyzed. As detailed below, this yielded a total of 67,908 regions of interest (ROI’s) from 793 brain volumes (Table 1).

**Table 1.**
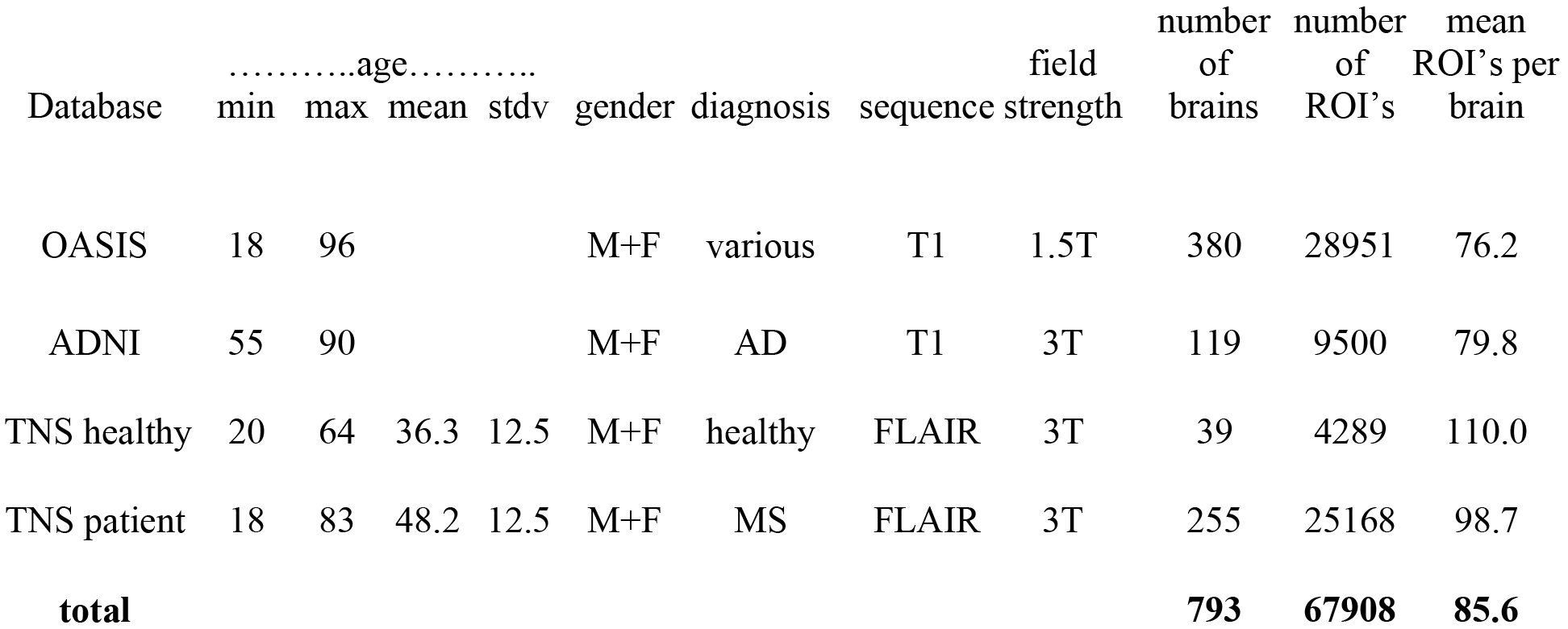
Characteristics of the MRI databases used in this study. AD: Alzheimer’s Disease. MS: multiple sclerosis.

Simulated T1 weighted MRI images were calculated using the BrainWeb simulator [2]–[4]. We used the system defaults, namely, a voxel size of 1.0 × 1.0 × 1.0 mm and spoiled FLASH with TR=18 ms, TE=10 ms, flip angle 30 deg. FLAIR MRI images with CSF suppression were not supported by the online BrainWeb simulator. To obtain these images, we modified the simulator code to use a T1 relaxation time of 4500 ms for CSF instead of the default value of 2569 ms, and a T2 relaxation time of 2300 ms for CSF rather than the default value of 329 [13] and simulated an inversion-recovery sequence with TR=11000 ms, TE=140 ms, TI =4600ms. These modifications required recompiling the simulator code locally, using Ubuntu 16.04.2. For both types of images, noise was simulated as additive Gaussian white noise, and its standard deviation was specified as a fraction of the most intense tissue value.

### 2.2 Processing Pipeline: Spectral Analysis

The processing pipeline was designed to extract individual regions of interest (ROI’s) that were fully contained within brain parenchyma in an automated, unbiased fashion (Figure 1). For each sagittal section, the skull was stripped using Freesurfer’s “watershed.” Then, using Matlab’s bwconvhull, the convex hull was determined, and the largest rectangle that fit inside this region was then found. Next, we randomly selected one 64 × 64 ROI from inside each rectangle (slices whose convex hulls were too small to contain a 64 × 64 rectangle were discarded). Optionally, to probe the possible effects of high-frequency artifacts in the reconstruction, the 64 × 64 ROI’s were downsampled to 32 × 32 ROI’s by averaging the intensities in 2×2 blocks. We present data from the full-resolution and the down-sampled analyses in parallel, to show that these artifacts have little effect on our findings.

**Figure 1.**
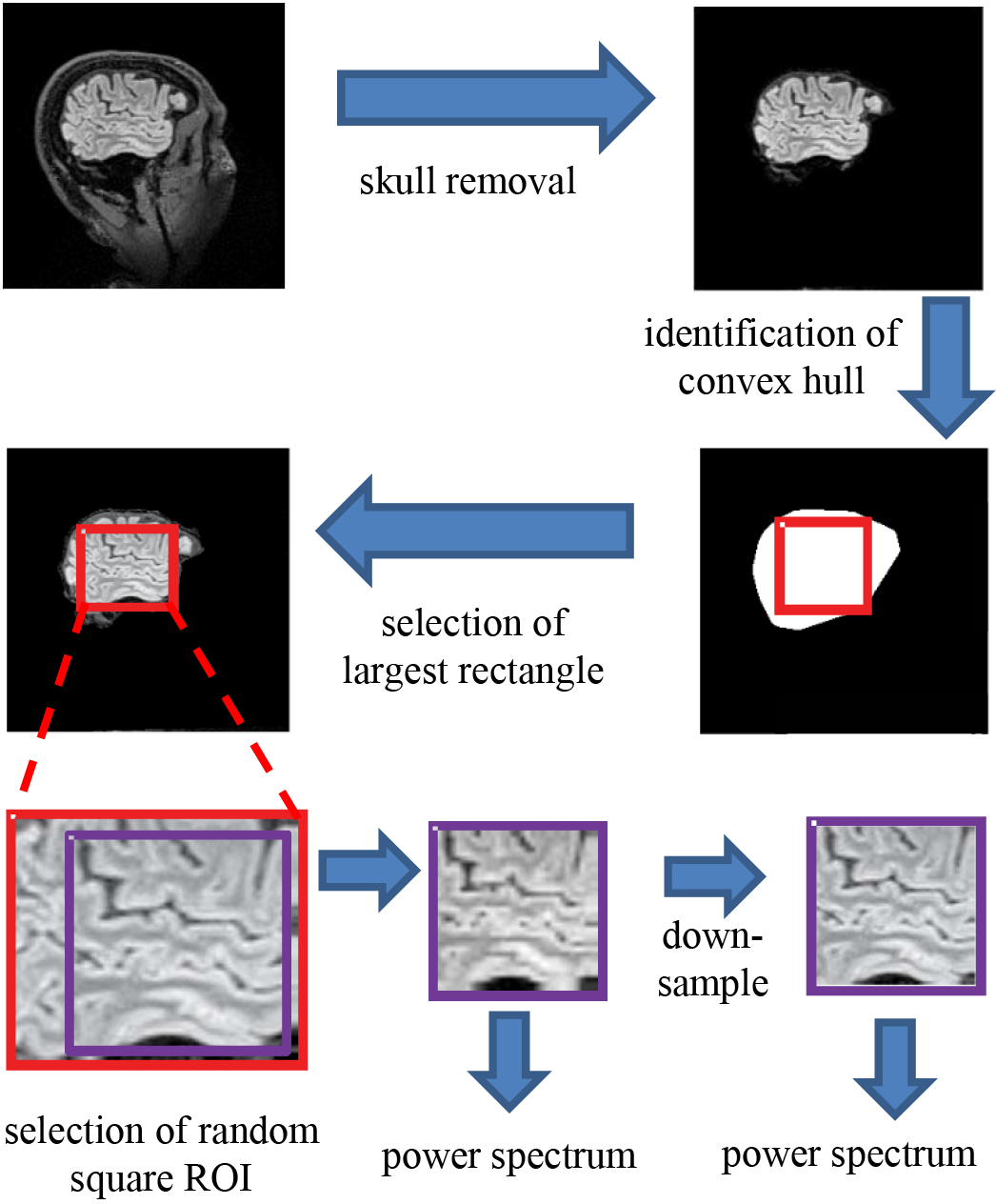
The processing pipeline for spectral analysis. Images of each sagittal plane were processed by (a) removal of the skull, (b) identification of the convex hull of the resulting image, (c) selection of the largest rectangle, and (d), selection of a random 64 × 64 pixel square ROI within this convex hull. The power spectrum was then computed from the mean of the squared amplitudes of the Fourier components obtained from these ROI’s. Optionally, the image was then downsampled by averaging within 2×2 blocks prior to Fourier transformation.

Spectra were computed by applying Matlab’s fft2 in each such region, and averaging the squared values across all slices from each database. Spectra were computed both with and without twofold padding. We focused on the frequency range from 2 to 16 cycles per ROI (1/32 to 0.25 cycles per pixel); in this range, padding (not used in the figures below) had minimal effect. For simplicity, we did not use windowing; the use of windows (e.g., a multitaper approach) would have little effect because the spectra are broadband.

To fit the power spectra to a power-law function, we regressed the logarithm of the spectral densities against the logarithm of spatial frequency (using Matlab’s regress), for spatial frequencies ranging from 2 cycles per ROI (1/32 cycles per pixel) to 10% below the Nyquist frequency.

### 2.3 Processing Pipeline: Local Image Statistics

Local image statistics characterize the features of images in small neighborhoods, and therefore they complement the statistical description provided by spectral analysis. We focus on the statistics that characterize the distribution of black-and-white colorings of 2×2 blocks of pixels. We choose this strategy for two reasons. First, they capture local features such as edges and corners. Second–critical to the goals of this study – – their distribution in natural images has been well-characterized[7], as has their salience to human observers [8]. The close match of these sets of findings [7] suggests that the human visual system is tuned to make efficient use of these natural image statistics, and motivated us to determine whether a similar match was present for MRI images.

Our procedure (Figure 2) is modeled after the processing pipeline used in that study, with parameters *R* (ROI size) and *N* (downsampling) of (*R*, *N*) = (64,1) and (*R*, *N*) = (32, 2). The larger (*R*, *N*) parameter values used by Hermundstad [7] for natural images (*R* × *N* > 64) could not be used, because of the limited number of voxels per plane in the MRI images.

**Figure 2.**
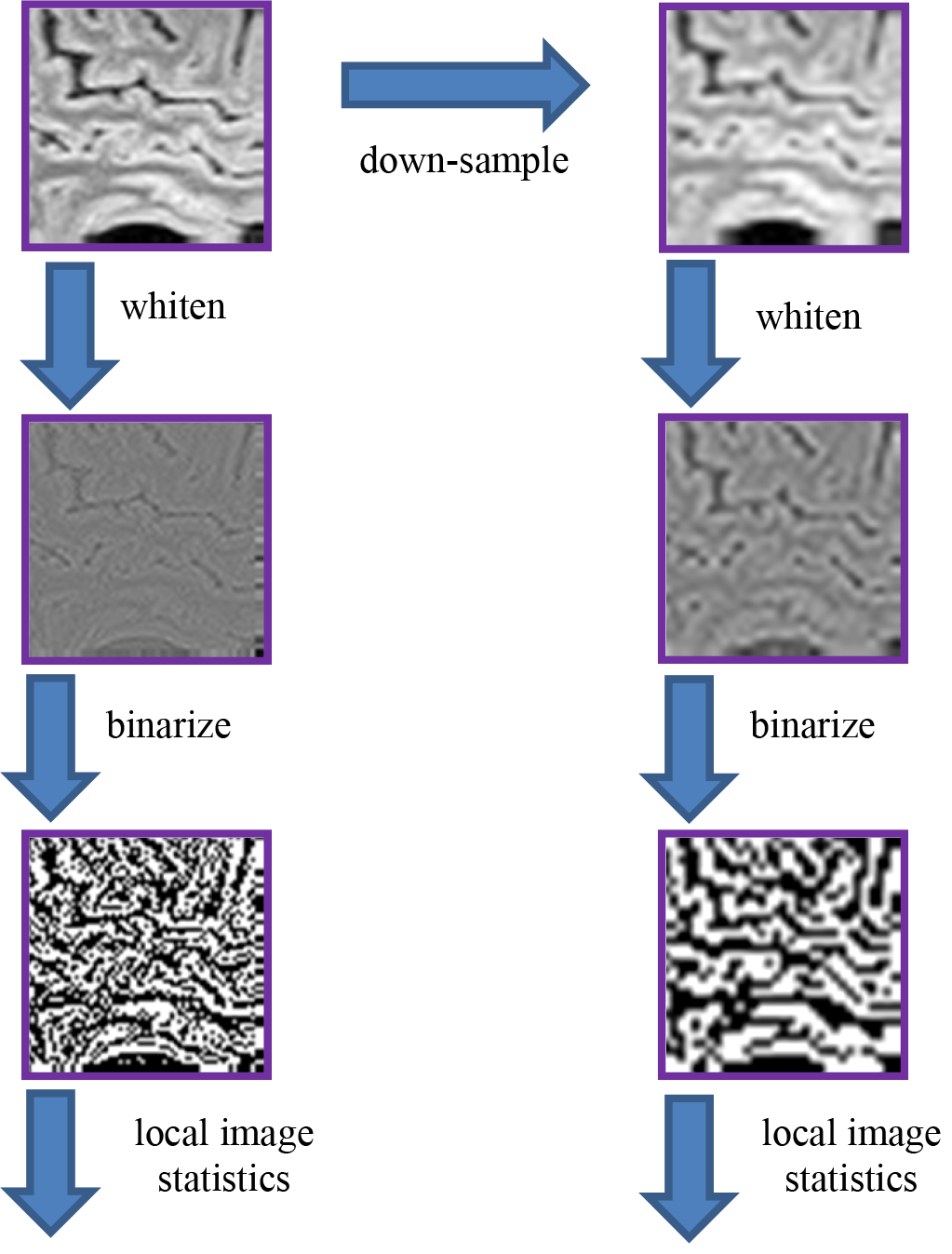
The processing pipeline for analysis of local image statistics. The ROI’s extracted for spectral analysis (Figure 1) were whitened by a linear filter whose amplitude is given by the square root of the power spectrum for that dataset. The resulting images were then binarized at the median value of the whitened images. Local image statistics were determined by tabulating the configurations of black and white checks within 2×2 neighborhoods.

For analysis of local image statistics, we began with the square ROI’s extracted above for spectral analysis (64 × 64 pixels, and also a parallel set downsampled to 32 × 32). Then, as in [7], we whitened these ROI’s by filtering them in the frequency domain, attenuating each Fourier component by the inverse square-root of the power spectrum. Then, these images were binarized at the median point, where the median was determined from all of the ROI’s drawn from the same brain. The binarized images were then parceled into 2×2 blocks, after removing a one-pixel border to avoid edge artifacts due to the whitening operation.

To describe the distribution of these blocks within an ROI, we proceeded as follows. Since each of the pixels in a block were either black or white, there are 16 (2^2×2^) different kinds of blocks. However, fewer than 16 degrees of freedom are required to describe them. These constraints arise because (a) the 16 probabilities add up to 1, and (b) the left two pixels of one block are also the right two pixels of the next (and similarly, for top and bottom). To take these constraints into account, we used the parameter set of [14], which uses a linear transformation to express the 16 raw block probabilities as 10 independent statistics.

These statistics, each of which ranges from −1 to 1, may be summarized as follows (see Figure 3). There are four two-point statistics, denoted *β*_|_, *β* _, *β*_\_, and *β*_/_. These describe the statistics of pairs of pixels, in directions corresponding to the subscript. For example, *β*_|_ is defined as the probability that two vertically-adjacent checks match, minus the probability that they do not match. So *β*_|_ = 1 means that all 2×1 (vertical) blocks are either both black or both white, while *β*_|_ = −1 means that all 2×1 pixel blocks contain mis-matching pixels. Similarly, *β* _ is the probability of matching vs. mismatching pixels in a1×2 (horizontal) block. *β*_\_ and *β*_/_ describe the probability that pixels which share a common corner either match or mismatch. Together, *β*_|_ and *β* _ will be referred to as the cardinal *β*’s; *β*_/_, and *β*_\_ will be referred to as the oblique *β*’s.

There are four three-point statistics, denoted *θ*_⌈_, *θ*_⌉_, *θ*_⌋_ and *θ*_⌊_.These coordinates indicate the probability of that L-shaped regions containing an even or an odd number of white pixels. *θ* = 1 means that there is always an odd number of white pixels in such a region, and *θ* = −1 means that there is always an even number of white pixels in the such a region, i.e., an odd number of black pixels. As shown in Figure 3, high or low *θ* values create regions with triangular shapes.

**Figure 3.**
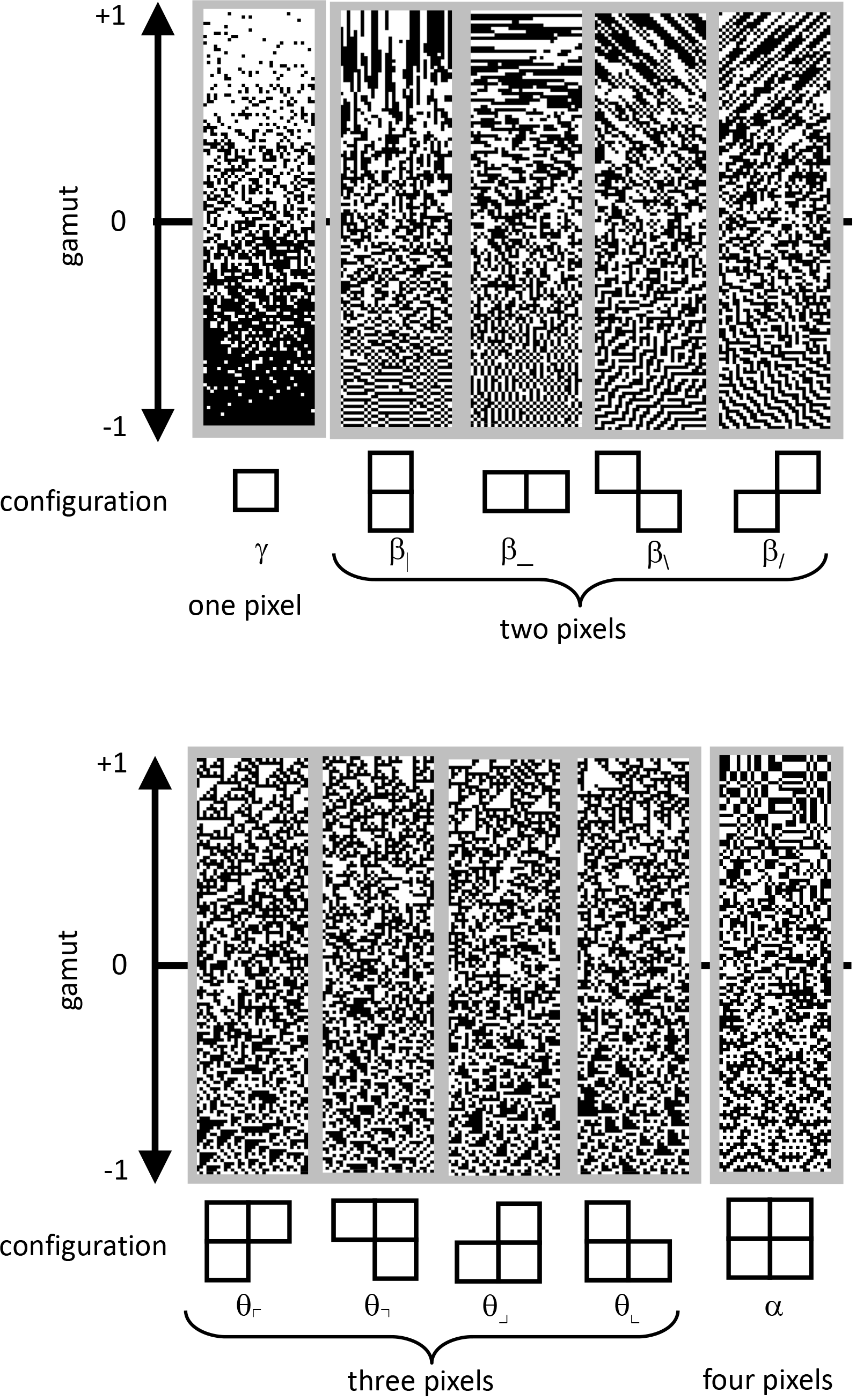
The parameterization of local image statistics used in this analysis. Each local statistic (*γ*, *β*_|_, *β*___, *β*_\_, *θ*_⌈_, *θ*_⌉_, *θ*_⌋_, *θ*_⌊_ and *α*) describes a specific kind of correlation between one or more pixels that form a template within a 2×2 neighborhood. Values of each statistic range from −1 to +1, with 0 indicating randomness. The strip above each template shows this gamut. Statistics are defined as follows. The one-point statistic *γ* is the difference between the probability that a check is white, vs. black: +1 means all white, −1 means all black. The two-point statistics *β*_|_, *β*___, *β*_\_ and *β*_/_ specify the probability that two nearby pixels match, minus the probability that they do not match: +1 means that they always match; −1 means that they always mismatch. The three-point statistics *θ*_⌈_,*θ*_⌉_,*θ*_⌋_ and *θ*_⌊_specify the difference between the fraction of L-shaped regions that contain an odd number of white pixels, and the fraction containing an even number: +1 produces an excess of white regions, while −1 produces an excess of black regions. The four-point statistic *α* specifies the difference between the probability that the number of white pixels in a 2×2 block is even, vs. odd: +1 means that 2×2 blocks always contain an even number of white pixels, −1 means that 2×2 blocks always contain an odd number of white pixels. Since the present analysis was carried out after binarization of at the median, *γ* was always close to zero, and not analyzed; it is shown for completeness. Adapted from Figure 1 of [8], with permission of the copyright holder, Elsevier B.V.

The four-point statistic *α* indicates the probability that the parity of the number of white pixels in a 2×2 block is even: *α* = 1 means that 2×2 blocks always contain an even number of white pixels and *α* = −1 means that 2×2 blocks always contain an odd number of white pixels.

Finally, there is a one-point statistic, γ, which is defined as the probability of a white check minus
the probability of a black check, and indicates the overall intensity. However, since binarization was carried out at the median, this statistic was always close to zero, and not further analyzed.

Together, the ten quantities {*β*_|_, *β*_, *β*_\_, *β*_/_, *θ*_⌈_, *θ*_⌉_, *θ*_⌋_, *θ*_⌊_, *α*, *γ*} are independent parameters that
specify the probabilities of the 2×2 blocks. We determined these quantities for each ROI, and then examined their means, variances, and covariances across the ROI’s within each database. To determine confidence limits on these estimates, we used a bootstrap procedure (500 resamplings). Resamplings were done with replacement of each slice. Each time a slice was drawn, a new square ROI was randomly chosen from the maximal rectangle within its convex hull.

## 3 Results

We characterize the statistics of MRI images in two complementary ways. We begin with the power spectrum, as it is a standard approach that captures global correlations. We then examine local image statistics, which are independent of the power spectra, and capture the features contained in small neighborhoods of pixels.

### 3.1 Spectral analysis

Figure 4 shows the spatial power spectra of images in the T1 weighted databases, plotted as a function of the magnitude of spatial frequency, along with the corresponding quantities computed from simulated T1 weighted data. For each database, there is a broad frequency range in which the power spectrum is approximately a power-law function of spatial frequency, i.e.,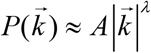, independent of whether images were analyzed at the full resolution or after downsampling. As is well-known, this power law relationship also holds for natural images, with the power-law exponent *λ* in the range −1.8 to −2.2 [15]–[17]. We therefore used this relationship to summarize and compare the spectra, and compare them with the spectra of natural images. Table 2 shows the exponent determined from each database, as well as from the simulated MRI data with various noise levels. For the T1 weighted images, this exponent was in the range −3.1 to −2.6 (see Table 2 for details and confidence limits), which is more negative than the values characteristic of natural images [15]–[17]. This means that, relative to natural images, MRI images have a greater amount of energy at low spatial frequencies. As the Figure 4 and Table 2 show, slopes more negative than −2 are also found in the simulated data, and slopes similar to the database images were found when a modest amount of noise (3%) was added.

**Figure 4.**
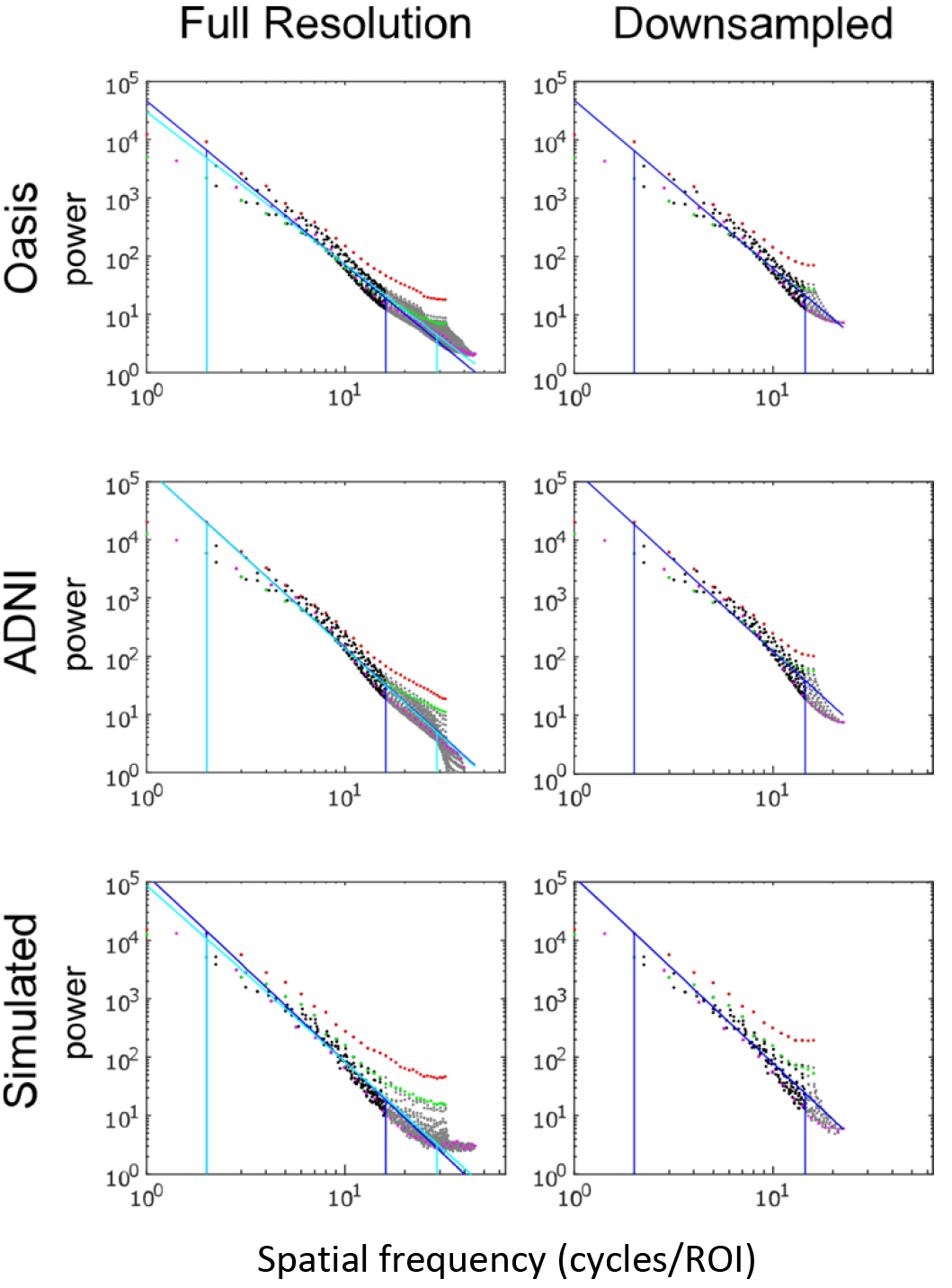
Power spectra of T1 MRI images, log-log plot. The simulated T1 images were constructed with 3% added noise; for further details, see text. Spectral estimates along the horizontal axis are shown in red, vertical in green, and oblique in magenta. The regression line, and the frequency range used to determine the regression, is shown for the full-resolution images in cyan, and for the 2×2 downsampled images in blue (with the latter superimposed on the full-resolution analysis for comparison purposes).

Beyond the spatial frequency range considered in the above analysis, MRI spectra become flatter, and are no longer well-fit by a single power-law function. This contrasts with the behavior of natural images, which have power-law behavior over a range of more than two decades [17]. However, because of the concern that these characteristics may reflect details of image reconstruction, smoothing, and noise, we do not further consider this behavior.

The two-dimensional power spectra (Figure 5) show that the MRI image spectra have significant anisotropies. In both T1 weighted databases, there is a prominent excess of power along the horizontal and vertical axes – amounting to a tenfold excess at the highest spatial frequencies. Such anisotropies are not typical of natural scenes, but have been noted in images containing manmade objects [15]. Away from the axes, the iso-intensity contours show a mild deviation from circularity, indicating a relative decrease of power in the oblique directions. Simulated T1weighted images share these features: they also have an excess of power along the horizontal and vertical axes, to a similar extent as in the MRI images drawn from the database.

**Figure 5.**
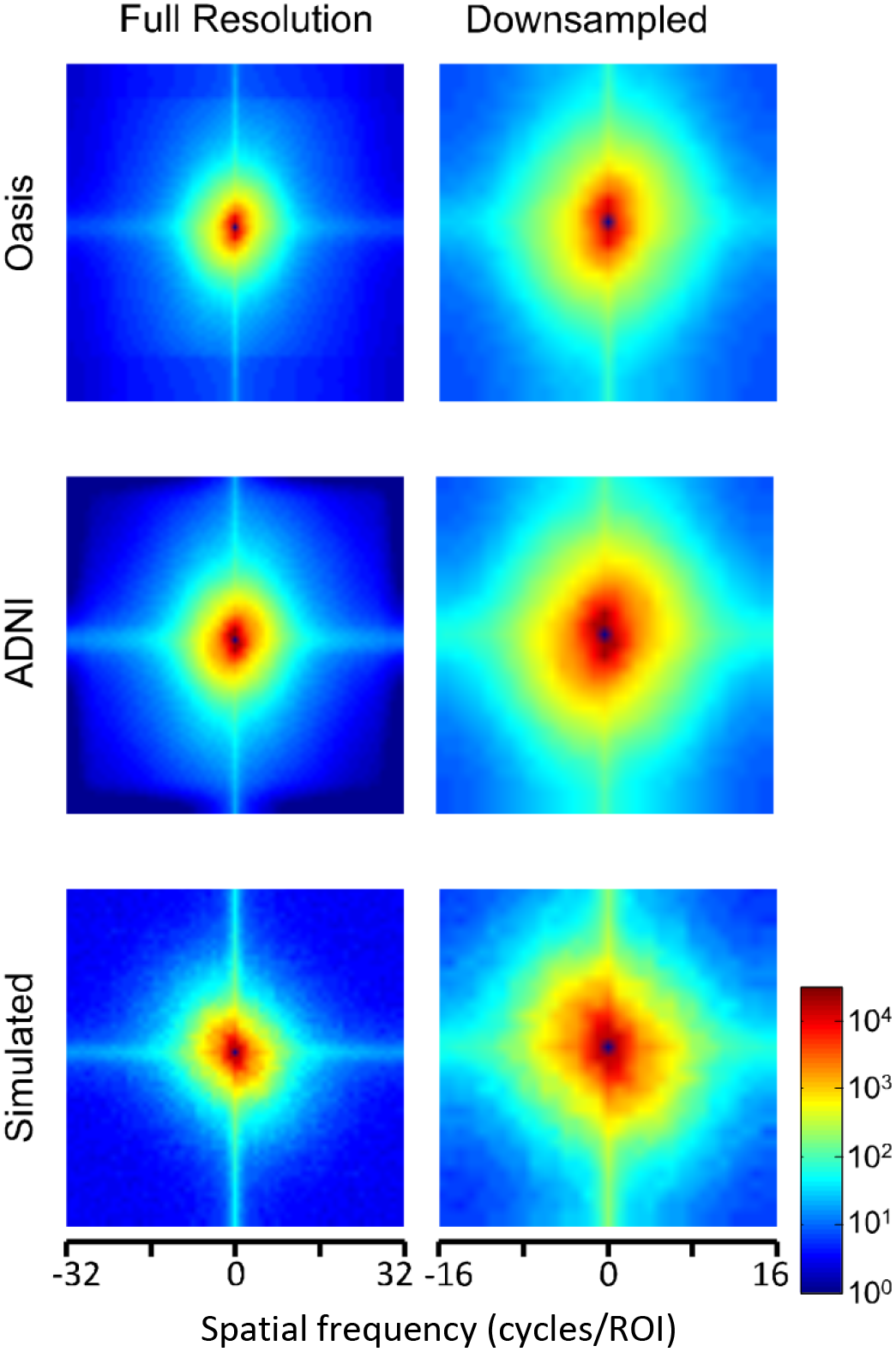
Power spectra of T1 MRI images (data of Figure 4), as a heatmap. Note that the color scale is logarithmic, and that the spatial frequency axes are expanded for the downsampled images (right column).

Power spectra of FLAIR images, taken from the TNS database, show these same characteristics (Figures 6 and 7, and Table 2): as was the case for the T1 images, spectral slopes were more negative than −2. The TNS database was subdivided into images from both healthy volunteers and patients; slopes were closely similar for the two subsets: −2.82 vs. −2.85 without downsampling, −2.25 vs. −2.21 with downsampling; overlapping confidence limits in both cases. Healthy volunteers and patients also both showed an excess of power, primarily along the vertical axis, of approximately a factor of three at the highest spatial frequencies. Simulated FLAIR images with 7% noise added (lower row of Figures 6 and 7) had a similar spectral slope as the MRI images. They also showed anisotropy to a similar degree, but (in contrast to the database images) a greater anisotropy along the horizontal axis than the vertical axis.

**Figure 6.**
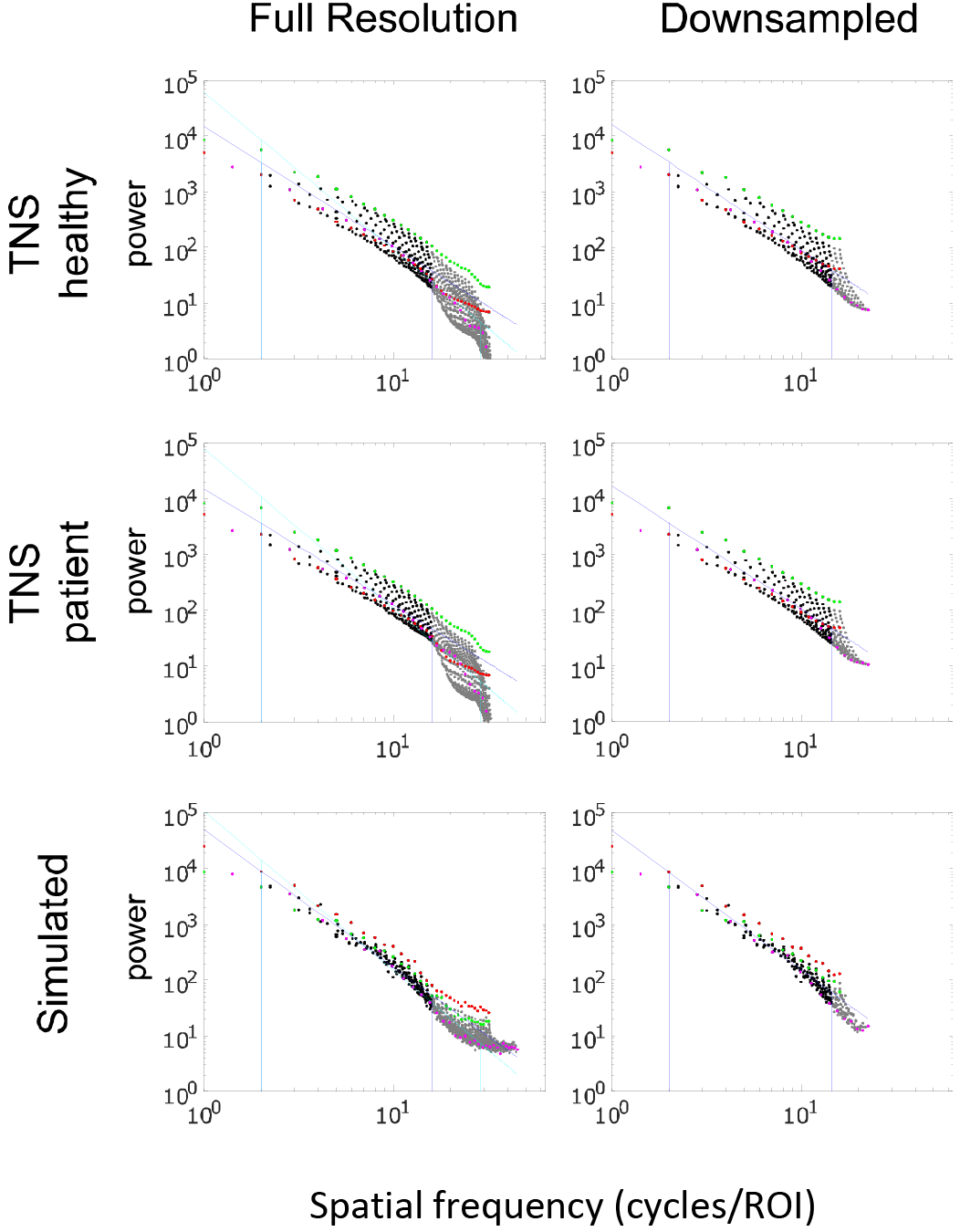
Power spectra of FLAIR MRI images, log-log plot. The simulated FLAIR images were constructed with 7% added noise; for further details, see text. Plotting conventions as in Figure 4.

**Figure 7.**
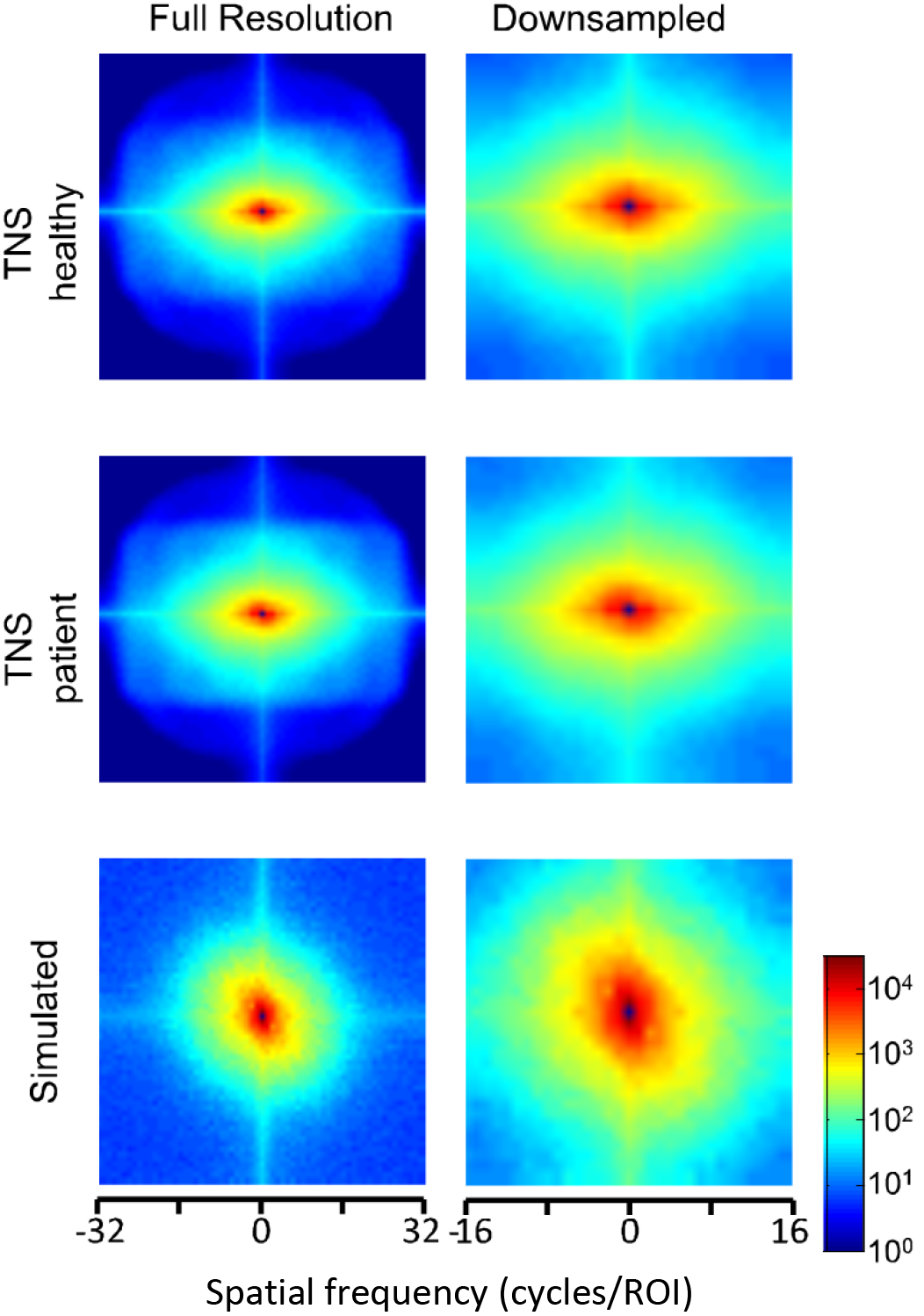
Power spectra of FLAIR MRI images (data of Figure 6), as a heatmap. Note that the color scale is logarithmic, and that the spatial frequency axes are expanded for the downsampled images (right column).

**Table 2.** Spectral slopes of MRI images and simulated images. Slopes were determined by regression of log(power) vs. log(spatial frequency) from 2 cycles per ROI to 90% of the Nyquist frequency (28.8 cycles per ROI for images processed at full resolution, 14.4 cycles per ROI for the downsampled images); this range is shown by the vertical blue and cyan lines in in Figures 4 and 6. Confidence limits (95%) were determined via t statistics, as implemented by Matlab’s regress.m routine.

|  | Full Resolution |  |  | Downsampled |  |  |
| --- | --- | --- | --- | --- | --- | --- |
|  | slope | confidence limits |  | slope | confidence limits |  |
| T1 images |  |  |  |  |  |  |
| OASIS | -2.61 | -2.67 | -2.56 | -2.88 | -3.00 | -2.75 |
| ADNI | -3.09 | -3.15 | -3.03 | -3.11 | -3.24 | -2.97 |
| simulated, no noise | -3.86 | -3.98 | -3.73 | -3.27 | -3.45 | -3.09 |
| simulated, 3% noise | -3.03 | -3.11 | -2.94 | -3.18 | -3.36 | -3.01 |
| simulated, 5% noise | -2.16 | -2.22 | -2.09 | -2.87 | -3.02 | -2.73 |
| simulated, 7% noise | -1.86 | -1.92 | -1.81 | -2.70 | -2.82 | -2.58 |
| FLAIR images |  |  |  |  |  |  |
| TNS healthy | -2.82 | -2.92 | -2.71 | -2.25 | -2.43 | -2.07 |
| TNS patient | -2.85 | -2.96 | -2.75 | -2.21 | -2.36 | -2.05 |
| simulated, no noise | -3.64 | -3.74 | -3.55 | -2.56 | -2.67 | -2.46 |
| simulated, 3% noise | -3.39 | -3.47 | -3.31 | -2.55 | -2.66 | -2.44 |
| simulated, 5% noise | -3.12 | -3.18 | -3.06 | -2.53 | -2.63 | -2.42 |
| simulated, 7% noise | -2.84 | -2.89 | -2.79 | -2.49 | -2.60 | -2.39 |

### 3.2 Local Image Statistics

We now characterize the local image statistics of MRI images. In brief (see Methods, Figure 2, and [7]), the approach consists of whitening the images followed by binarization, and then tabulating the configurations in 2×2 neighborhoods of pixels within each ROI. This tabulation is carried out in terms of a 9-parameter set of descriptors (see Figure 3), which are independent, and stratify the analysis into statistics involving two points(*β* _, *β*_|_, *β*_/_, and *β*_\_), three points (*θ*_⌈_,*θ*_⌉_, *θ*_⌋_, and *θ*_⌊_), and four points (the statistic *α*). Thus, the local statistical analysis is complementary to the characterization via power spectra - both because it focuses on the features in small neighborhoods of pixels, and also, because it is carried out after a whitening step that removes the overall correlations that shape the power spectrum. Because of this whitening step, any differences between the local image statistics of MRI images and those of natural images cannot be attributed to differences in their overall spatial frequency content.

We consider both the mean and standard deviations of these local statistics measured in each ROI. The mean values capture the average characteristics of all ROIs within a database, and thus focus on overall differences between the databases. The standard deviations describe the characteristics of the ROIs that distinguish them from each other, and thus focus on characteristics that are useful for analyzing individual MRI images. To enable a direct comparison to natural images, we accompany the analysis by a parallel analysis of natural images (recomputed from the raw data of [7], provided by Ann Hermundstad). Following Hermundstad [7], natural images were only analyzed after twofold downsampling (*N* = 2, *R* = 32), to avoid possible camera-related artifacts.

Figure 8 shows the mean values of the image statistics for images analyzed at full resolution (Panel A), and after twofold downsampling (Panel B). At both scales, there were substantial differences in database characteristics: For example, at native resolution, vertical two-point correlations (*β*_|_) were zero or positive for the T1 databases but negative for the FLAIR databases, while horizontal two-point correlations (*β*_) were negative for the T1 databases but positive for the FLAIR databases. Other differences were present for three- and four-point correlations at native resolution, and for many of the statistics after twofold downsampling. In the latter analysis, there were also differences between the two T1 databases across all local image statistics. For the simulated MRI images, many of the mean values differ from mean values obtained from actual MRI images, for both sequence types and at both scales. This complex pattern of variations is not surprising, as the means of these statistics will be affected by the MRI intensity histogram, which will vary across sequence types as a consequence of the physics of signal generation and, possibly, differences in the reconstruction process. Thus, the means of the image statistics are affected by many sequence-dependent factors in addition to brain morphology at the MRI scale. For completeness, Fig. 8B also shows the mean values of local image statistics obtained from natural images.

**Figure 8.**
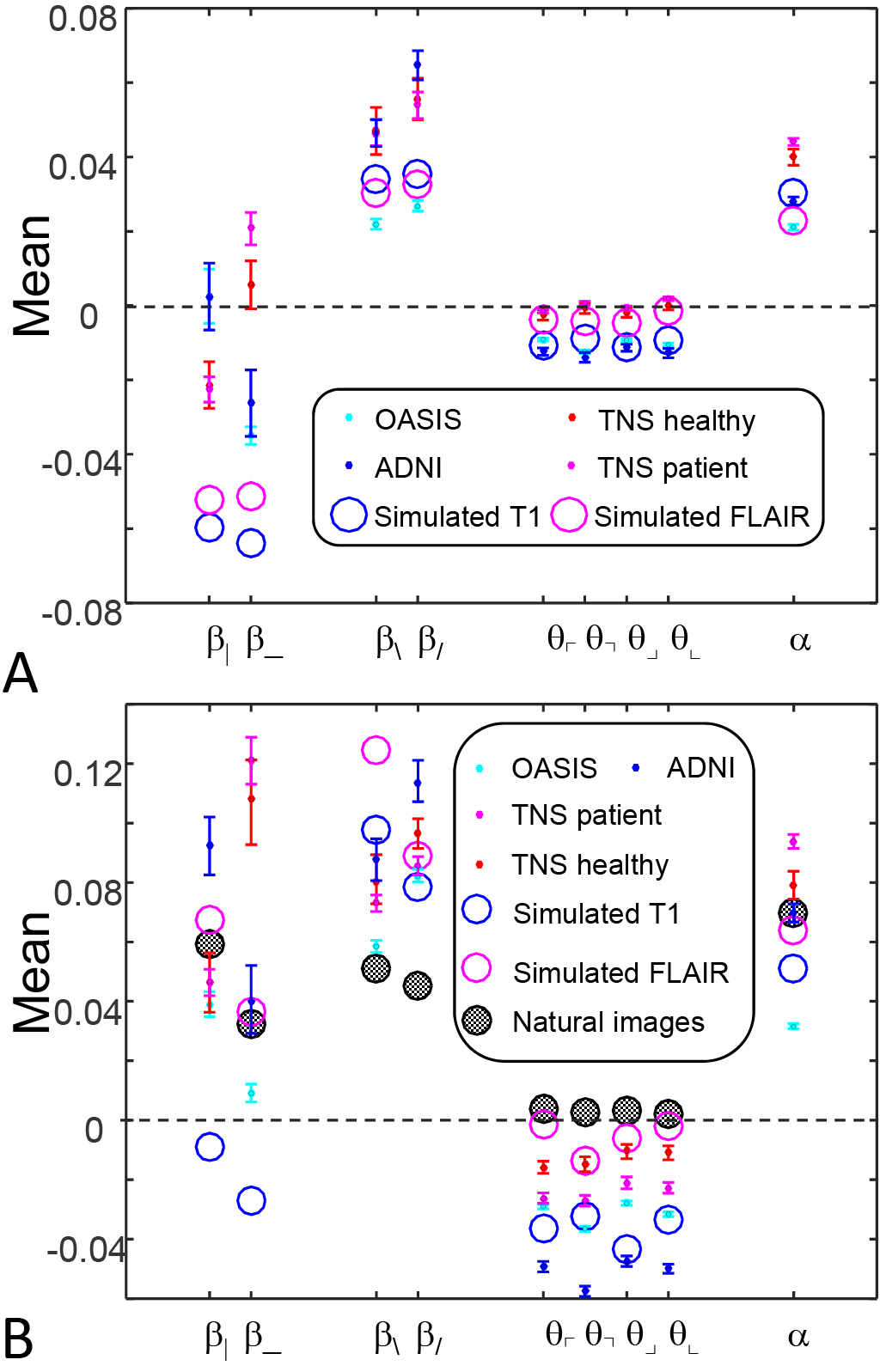
Means of local image statistics for real (solid symbols) and simulated (open symbol) MRI images. Panel A: full resolution ((*R*, *N*) × (64,1)); panel B: after 2×2 downsampling ((*R*, *N*) × (32, 2)). In B, values for the natural image database of [7] are shown (shaded symbols). For MRI images, error bars indicate 95% confidence intervals, computed via bootstrap.

In contrast, the standard deviations of these image statistics – which quantify their ability to distinguish one ROI from another -- show a simpler and more consistent pattern (Fig. 9). Two-point statistics in the vertical and horizontal directions have the largest standard deviations, followed by two point statistics in the oblique directions. Three- and four-point statistics have smaller but comparable standard deviations. This pattern is seen for images at full resolution (Fig. 9A) and after 2×2 downsampling (Fig. 9B), and also for the simulated T1 and FLAIR MRI images. The pattern is also consistent between images obtained from normal volunteers and patients within the TNS database.

While these statistical characteristics are common to images from all the MRI databases and the simulated images, they differ from natural images (Fig. 9B). Specifically, in comparison to natural images, horizontal and vertical two-point statistics are less variable across patches (*β*_|_ and *β*___ of Fig. 9B): in all cases, the standard deviations of the MRI statistics are below the values for natural images. The opposite pattern is present for three-point statistics (the *θ*’s of Fig. 9B): standard deviations for the MRI statistics are above the corresponding values for natural images. These differences are consistent across real and
simulated MRI images, and are substantially in excess of the confidence limits. That is, two-point correlations are relatively more stereotyped for MRI images than for natural images, while three-point correlations are relatively more stereotyped for natural images than for MRI. Consequently, the relative importance of two- and three-point local image statistics for distinguishing among patches of MRI images differs from their relative importance in natural images.

**Figure 9.**
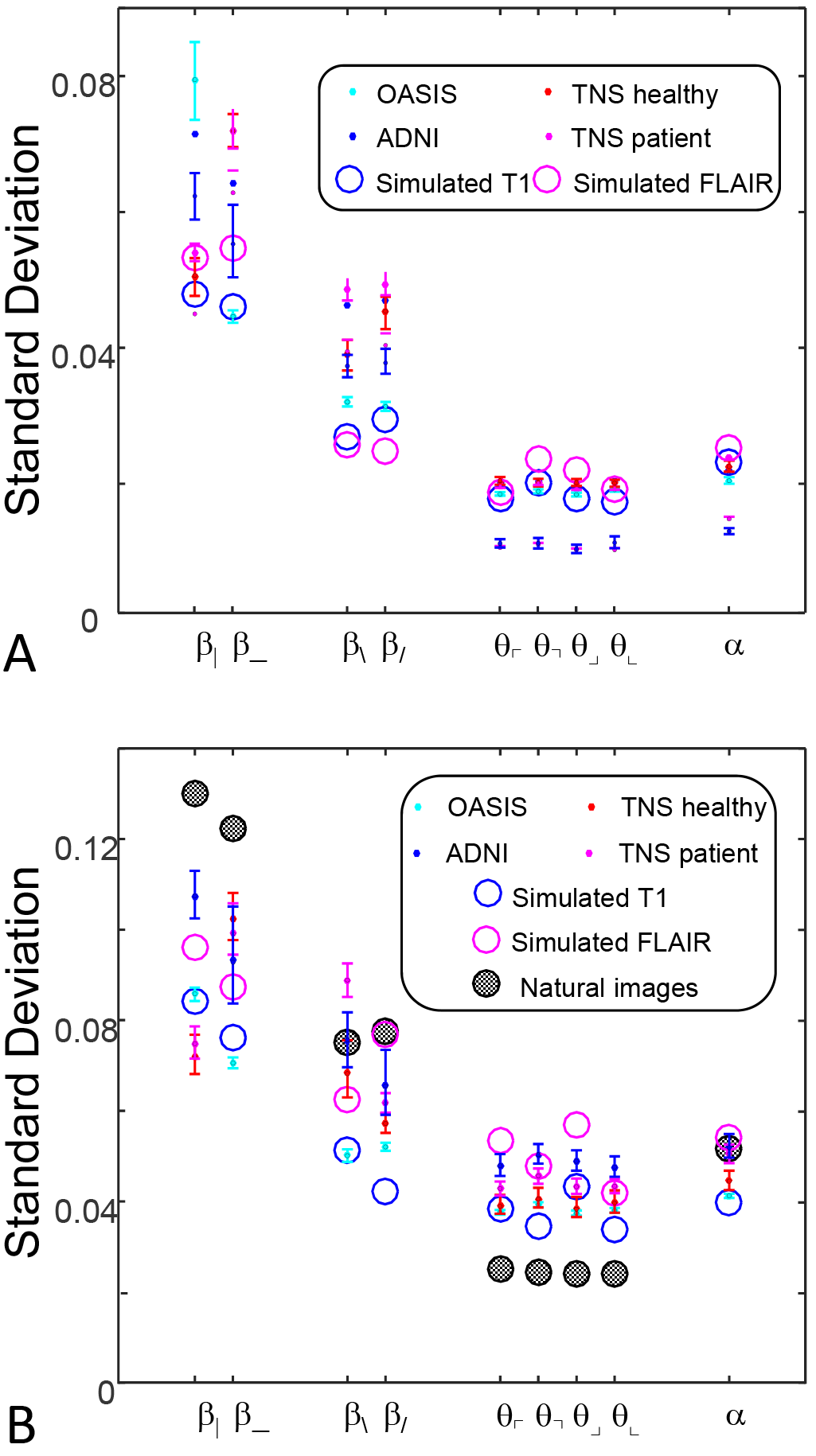
Standard deviations of local image statistics for real (solid symbols) and simulated (open symbol) MRI images. Panel A: full resolution ((*R*, *N*) × (64,1)); panel B: after 2×2 downsampling ((*R*, *N*) × (32, 2)). Other details as in Figure 8.

For completeness, Figure 10 shows the covariances of the image statistics, normalized to their corresponding standard deviations described above. With this normalization, there are few differences between the MRI images, or between them and the corresponding statistics of natural images.

**Figure 10.**
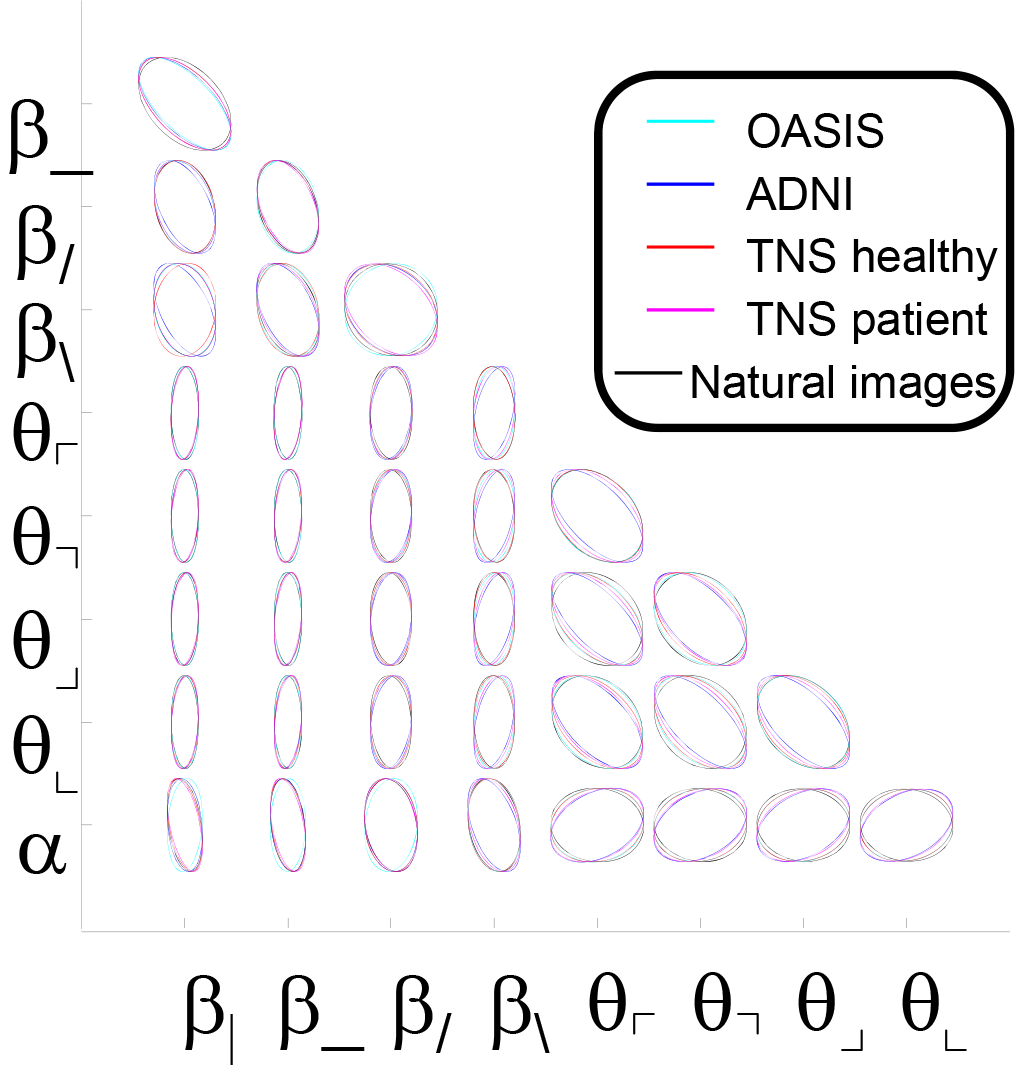
Covariances of local image statistics. Covariances for each pair of local image statistics are represented by their precision matrix, rendered as an ellipse scaled to maximally fill each portion of the grid. Ellipses that are nearly circular indicate that the statistics are independent. Elongation of the ellipse along an axis indicates that the corresponding linear combination of statistics is “precise”, i.e., has a value that varies little across ROI’s.

## 4 Discussion

Drawing inferences from medical images is the end result of a processing pipeline, whose stages include not only the process of image formation but also visual analysis by human observers. The ability of humans to interpret images is the result of evolutionary pressures in the natural environment. It is therefore shaped by, and tuned to, the statistical properties of natural images, including both their global correlation properties as captured by their power spectrum [16], and their local image statistics [7], [8]. These statistical properties arise from the physics of image formation and the characteristics of the natural environment. Medical images are formed by different physical processes – and this raises the possibility that medical images form a category of images whose statistics differ from those that our visual systems are adapted to process.

We found that brain MRI’s are a category of images whose statistics differ from those of natural images in several ways. With regard to global correlation properties, the spectral slope of MRI images is in the range of −2.2 to −3, steeper than the slope of −1.8 to −2.2 characteristic of natural images [15]–[17]. A spectral slope of −2 – the middle of the range found for natural images - corresponds to scale invariance, which follows from the notion that an observer’s viewpoint is chosen at random with respect to the environment. MRI images do not have this characteristic, and are more heavily weighted towards low spatial frequencies (Figs. 4-7).

The statistics of local features of natural images are less well-studied, but these too have consistent characteristics to which the human visual system is tuned. We focus on the local features that are important in distinguishing one image patch from another, i.e., the extent to which image statistics vary across ROI’s (Fig. 9) For natural images, two-point correlations are the most informative; correlations in cardinal directions (vertical and horizontal) are substantially more important than correlations in oblique directions; this corresponds to their perceptual salience [7], [8]. In contrast, for MRI images, cardinal and oblique correlations are more nearly equal in variance, and hence, in importance for distinguishing among patches (Fig. 9B). With regard to higher-order statistics, three-point statistics are less important than four-point statistics in natural images, but these are comparable for MRI. (Fig. 9B). Note that the local statistical features are analyzed after a whitening step, and thus, are independent of the differences in spectral characteristics.

Each MRI sequence has its own tradeoffs between bandwidth, noise, contrast and artifacts, and, a priori, any of these could have had a role in determining the statistics of resulting images. Interestingly, we found consistency across three databases representing two sequence types (T1 weighted and pre contrast FLAIR) as well as simulated MRI images. Based on this consistency, we infer that these statistical characteristics presented here are not due to the physics specific to each kind of MRI sequence, or filtering and artifacts that might arise in the process of data acquisition or reconstruction, but rather reflect the 3D geometry of the brain and its sectioning into 2D slices at the resolution of MRI. That our results are consistent not only across two MRI modalities and but also across simulations suggests that our results are not driven by steps in the acquisition process. This logic also suggests that other medical image categories – such as transmission X-rays, light micrographs, and electron micrographs – will also have their own characteristic statistical fingerprints.

These observations have a number of implications. First, the fact that human brains are attuned evolutionarily to process natural images whose statistics differ from that of brain MRIs suggests that the human visual system is not ideally matched to the challenges of reading MR exams. For example, three point and four-point statistics are similarly important for distinguishing the among MRI regions, while in natural images, four-point statistics are substantially more important (Figure 9B). Human observers [7],[8] allocate their computational resources in a way that is matched to the characteristics of natural images: they are more sensitive to four-point statistics than three-point statistics. However, there are modest inter individual differences in sensitivity to local image statistics [8], [14]; one may speculate that the individuals with relatively greater sensitivity to the third-order statistics that are informative in MRI images will more readily develop neuroradiographic expertise. Conversely, the development of such expertise through training may be associated with increased visual sensitivity to the characteristic statistical features of MRI. Of note, visual expertise is well-recognized in many domains, including identifying distinguishing features in plain radiographs [18], [19], cytological images [19], and fingerprint identification [20]. Here we raise the possibility that such expertise is related to distinctive statistical characteristics of the image set.

These observations also may be useful in developing improved methods of MRI analysis. We recognize that it is very nontrivial to go from a characterization of image statistics to an image-processing algorithm that exploits these characteristics – but several possibilities naturally arise. (i) Because MRI images have a distinctive and stereotyped statistical structure, deviations of an acquired image from these norms is a potential strategy for artifact detection and quality control. (ii) Super-resolution methods are applied to enhance a reconstructed image, but, because of the ill-posed nature of the problem, regularization methods are typically required [21]. Incorporating the distinctive statistical characteristics of brain MRI into a prior may be a way to develop more effective statistical regularization methods. In these two applications, a cost function for image patches, parameterized by their local image statistics, could be constructed as a log likelihood based on the multidimensional Gaussian that yields the means, variances, and covariances of these statistics in MRI datasets.

Finally, (iii) it may be possible to improve the interpretability of MRI images by local transformations that make their textural features more visible to human observers. In this regard, two recent studies are notable: a study that demonstrated the utility of second-order statistics in distinguishing T2 MRI images of normals and patients with Alzheimer’s disease [22](though local and long-range statistics were not distinguished in that study), and a study that demonstrated superiority of machine analysis of MRI images to human analysis in distinguishing tumor from radionecrosis [23][24]. While neither of these studies made use of the specific image statistics studied here, they support the notion that textural features other than those computed by the human visual system are important for MRI interpretation.

## 5 Acknowledgements

We thank Mohammad Ali (at Carbonite), Jim Chen (at Kaiser Permanente), Jason Econome (at Stuyvesant High School), Ramin Zabih, and Amy Kuceyeski (both at Cornell) for their insightful comments. We also thank Daniel S. Reich and the NINDS for access to the TNS database. JV is supported by NIH R01 E07977 from the NEI. We also acknowledge NIH grants P50 AG05681, P01 AG03991, R01 AG021910, P20 MH071616, U24 RR021382 for support of OASIS.

Data collection and sharing for this project was funded by the Alzheimer’s Disease Neuroimaging Initiative (ADNI) (National Institutes of Health Grant U01 AG024904) and DOD ADNI (Department of Defense award number W81XWH-12-2-0012). ADNI is funded by the National Institute on Aging, the National Institute of Biomedical Imaging and Bioengineering, and through generous contributions from the following: AbbVie, Alzheimer’s Association; Alzheimer’s Drug Discovery Foundation; Araclon Biotech; BioClinica, Inc.; Biogen; Bristol-Myers Squibb Company; CereSpir, Inc.; Cogstate; Eisai Inc.; Elan Pharmaceuticals, Inc.; Eli Lilly and Company; EuroImmun; F. Hoffmann-La Roche Ltd and its affiliated company Genentech, Inc.; Fujirebio; GE Healthcare; IXICO Ltd.; Janssen Alzheimer Immunotherapy Research & Development, LLC.; Johnson & Johnson Pharmaceutical Research & Development LLC.; Lumosity; Lundbeck; Merck & Co., Inc.; Meso Scale Diagnostics, LLC.; NeuroRx Research; Neurotrack Technologies; Novartis Pharmaceuticals Corporation; Pfizer Inc.; Piramal Imaging; Servier; Takeda Pharmaceutical Company; and Transition Therapeutics. The Canadian Institutes of Health Research is providing funds to support ADNI clinical sites in Canada. Private sector contributions are facilitated by the Foundation for the National Institutes of Health (www.fnih.org). The grantee organization is the Northern California Institute for Research and Education, and the study is coordinated by the Alzheimer’s Therapeutic Research Institute at the University of Southern California. ADNI data are disseminated by the Laboratory for Neuro Imaging at the University of Southern California.

